# Multi-omic mitoprotease profiling defines a role for Oct1p in coenzyme Q production

**DOI:** 10.1101/155044

**Authors:** Mike T. Veling, Andrew G. Reidenbach, Elyse C. Freiberger, Nicholas W. Kwiecien, Paul D. Hutchins, Michael J. Drahnak, Adam Jochem, Arne Ulbrich, Matthew J.P. Rush, Joshua J. Coon, David J. Pagliarini

**Affiliations:** Morgridge Institute for Research, Madison, WI 53715, USA; Department of Biochemistry, University of Wisconsin–Madison, Madison, WI 53706, USA; Genome Center of Wisconsin, Madison, WI 53706, USA; Department of Chemistry, University of Wisconsin–Madison, Madison, WI 53706, USA; Department of Biomolecular Chemistry, University of Wisconsin–Madison, Madison, WI 53706, USA

**Keywords:** Mitoproteases, protease, oligopeptidase, Oct1p, MIPEP, coenzyme Q, Coq5p, mitochondria, multi-omic, transomic

## Abstract

Mitoproteases are becoming recognized as key regulators of diverse mitochondrial functions, although their direct substrates are often difficult to discern. Through multi-omic profiling of diverse *Saccharomyces cerevisiae* mitoprotease deletion strains, we predicted numerous associations between mitoproteases and distinct mitochondrial processes. These include a strong association between the mitochondrial matrix octapeptidase Oct1p and coenzyme Q (CoQ) biosynthesis—a pathway essential for mitochondrial respiration. Through Edman sequencing, and *in vitro* and *in vivo* biochemistry, we demonstrated that Oct1p directly processes the N-terminus of the CoQ-related methyltransferase, Coq5p, which markedly improves its stability. A single mutation to the Oct1p recognition motif in Coq5p disrupted its processing *in vivo*, leading to CoQ deficiency and respiratory incompetence. This work defines the Oct1p processing of Coq5p as an essential post-translational event for proper CoQ production. Our custom data visualization tool enables efficient exploration of mitoprotease profiles that can serve as the basis for future mechanistic investigations.

## INTRODUCTION

Mitoproteases—a collection of proteases and peptidases that reside in or translocate to mitochondria (Quiros et al.,2015)—are involved in many aspects of mitochondrial function. Once considered important merely for the removal of damaged proteins, it is now appreciated that mitoproteases are involved in an intricate protein quality control system that maintains mitochondrial proteostasis by ensuring proper protein import and processing (Ieva et al., 2013), influencing the half-lives of key regulatory proteins (Vögtle et al., 2011), and activating/deactivating proteins essential for core mitochondrial activities in a highly specific and regulated manner (Anand et al., 2014; Konig et al., 2016; Koppen and Langer, 2007). Consistently, mitoproteases are connected to the aging process, and their disruption underlies many pathological conditions, including cancer (Fukuda et al., 2007; Quiros et al., 2014), metabolic syndrome (Almontashiri et al., 2014; Civitarese et al., 2010), and neurodegenerative disorders (Di Bella et al., 2010; Strauss et al., 2005; Wai et al., 2015)

Given the increased appreciation for the importance of mitoproteases in basic mitochondrial biology and human health, multiple recent investigations have attempted to define their direct modes of action and breadth of activity. These studies employed diverse methodologies, including pulse-chase experiments (Christiano et al., 2014), proteomics analyses (Avci et al., 2014), and the development of “trap” mutants (Grimsrud et al., 2012; Trentini et al., 2016), among others. Despite the relative success of these approaches, many mitoproteases lack more than a few verified substrates. This stems from biochemical challenges, such as difficulties in purifying proteins for *in vitro* analyses and indirectly capturing protease-substrate interactions, but also from a general lack of knowledge about what pathways are associated with each protease. Broader analyses that succeed in associating mitoproteases within specific mitochondrial pathways could facilitate the discovery of new substrates.

Recently, we devised a mass spectrometry-based multi-omics approach designed to predict functions for <u>m</u>itochondrial uncharacterized (<u>x</u>) <u>p</u>roteins (MXPs) (Stefely et al., 2016).This approach was built on the underlying proposition that MXPs could be linked to proteins of known pathways and processes by virtue of the whole-cell multi-omic signatures that resulted from their respective gene deletions. To broadly explore connections between diverse mitoproteases and specific mitochondrial functions, we performed a similar multi-omic analysis of 20 yeast strains including knockout strains for nearly all known mitoproteases. These measurements were made using both gas and liquid chromatography coupled with high resolution Orbitrap mass analysis (Hebert et al., 2014b; Peterson et al., 2014a; Peterson et al., 2014b; Peterson et al., 2010; Richards et al., 2015a). From 360 individual LC and GC-MS experiments, we detected and quantified 4,031 proteins, 486 metabolites, and 52 lipids. Analysis of these data suggest numerous associations between individual mitoproteases and distinct mitochondrial processes, including protein import, complex assembly, metal ion homeostasis, cardiolipin metabolism, and metabolite transport.

These analyses revealed a particularly strong protein-, lipid-, and metabolite-based association between the mitoprotease Oct1p and CoQ. Oct1p is an octapeptidyl aminopeptidase located in the mitochondrial matrix that cleaves eight amino acids off the N-termini of select proteins following cleavage of their mitochondrial localization sequences (MLS) by the mitochondrial processing protease (MPP) (Isaya et al., 1991). To date, 14 substrates have been reported for Oct1p and its disease-related metazoan ortholog, MIPEP, but it has been speculated that multiple others might exist (Eldomery et al., 2016; Vögtle et al.,2011). The full functional relevance of Oct1p processing is debated, but it often it often serves to increase protein stability by establishing a new N-terminal residue that is more favorable according to N-end rules (Varshavsky, 2011).

To search for CoQ-associated substrates of Oct1p, we analyzed the N-termini of isolated CoQ-related proteins from wild type (WT) and Δ*oct1 Saccharomyces cerevisiae* using Edman sequencing. This analysis identified Coq5p as a direct Oct1p substrate, which we confirmed using *in vitro* protease assays. Coq5p and its human ortholog COQ5 are core members of complex Q, the CoQ biosynthesis complex (Floyd et al., 2016; Nguyen et al., 2014). Aberrant expression of Coq5p can inhibit cell growth, suggesting that Coq5p expression *in vivo* might be regulated at multiple levels (Lapointe et al., 2017). We further demonstrated that disrupted Oct1p processing *in vivo* causes a marked decrease in Coq5p stability, leading to CoQ deficiency and respiratory incompetence.

Overall, our work implicates a mitoprotease in a requisite post-translational processing step for proper CoQ biosynthesis. Beyond this, our multi-omic analyses provide many additional insights into the substrates and activities of mitoproteases. These large-scale data can be freely accessed and analyzed using our interactive data analysis tool at: http://mitoproteaseprofiling.com.

## RESULTS

### Multi-omic profiling connects mitoproteases to diverse mitochondrial processes

Hundreds of mitochondrial proteins have no or incompletely established functions (termed MXPs), and many mitochondrial pathways involve processes not currently assigned to a specific protein. To further investigate the potential roles for mitoproteases in mitochondrial biology, we performed a multi-omic analysis modeled on the experimental design of our multi-omic “Y3K” investigation (Stefely et al., 2016). To do so, we grew wild-type (WT) yeast along with 19 individual yeast deletion strains (14 intrinsic mitoproteases, three extramitochondrial proteases, one pseudoprotease, and Mcx1p, a homolog of the ClpX ATPase subunit of the ClpXP system) (Table S1) (Rottgers et al.,2002; Tzagoloff et al., 1986; van Dyck et al., 1998).Each strain was grown in biological triplicate under both standard fermentation culture conditions and optimized respiration culture conditions (Figure 1A, and Table S1), and analyzed using three separate high-resolution mass spectrometry (MS)-based proteomic, metabolomic, and lipidomic techniques (Figure 1B). This set of strains includes knockouts of all known yeast mitoproteases except Δ*mas1* and Δ*mas2*, which are inviable (Jensen and Yaffe, 1988; Witte et al., 1988).Altogether, these analyses quantified 4031 proteins, 486 metabolites, and 52 lipids across 360 individual MS experiments (Figure 1B, Table S2).

**Figure 1.**
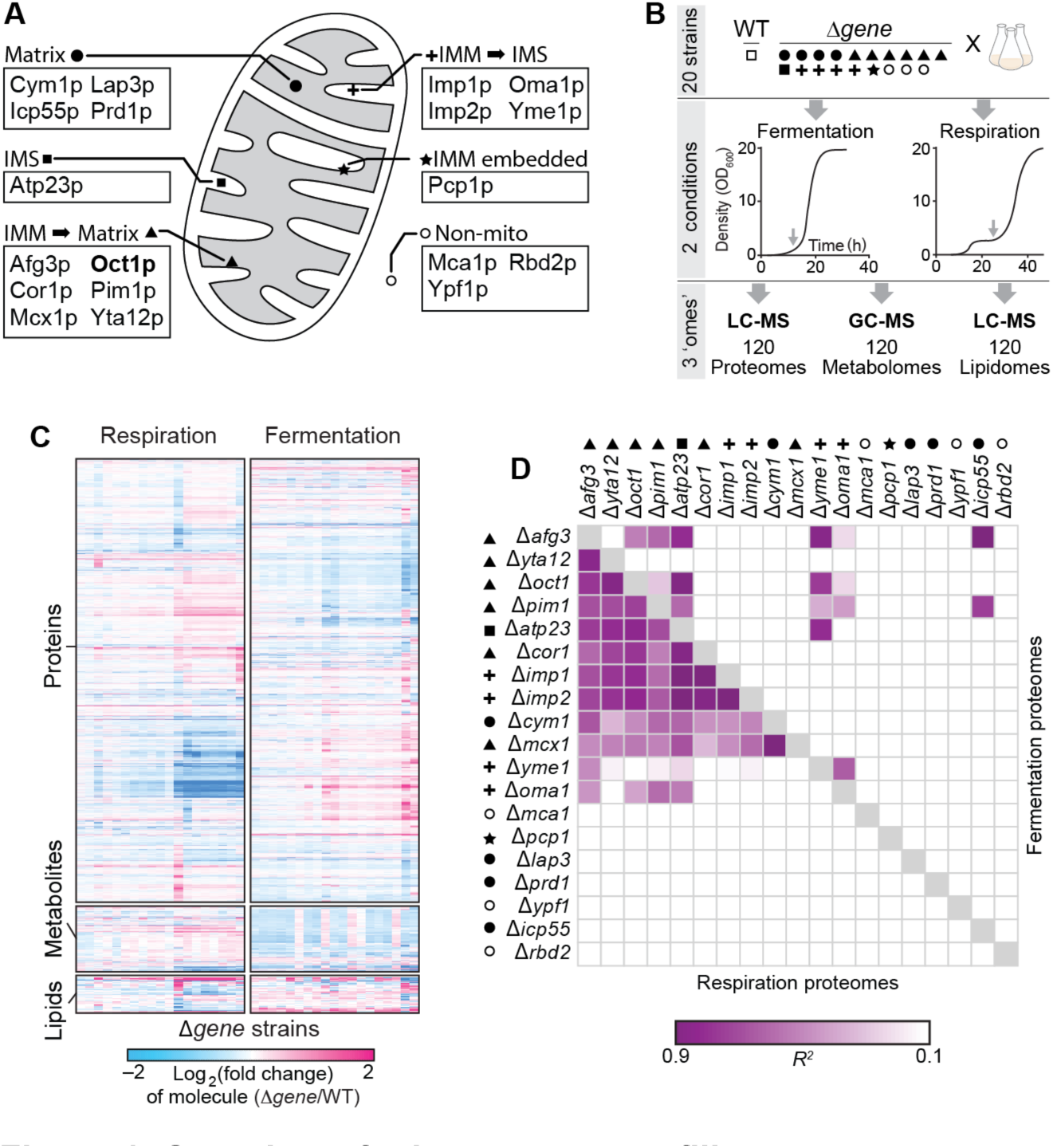
Overview of mitoprotease profiling. (A) Proteins encoded by the individual genes knocked out of the 19 yeast strains investigated in this study,shown in the context of their mitochondrial localization. (B) Overview of experimental design and data collected. Shapes indicate protein localization as shown in A. Representative growth curves are shown to indicate the yeast growth stage at time of collection. (C) Hierarchical clusters of Δ*gene* strains in respiration and fermentation across the proteome metabolome, and lipidome. Clustering was based on relative abundances compared to WT for significantly changing proteins as quantified by MS (mean, *n* = 3, *P* < 0.05, two-sided Student’s *t*-test). Strains are clustered based on respiration proteome correlations for all maps. See Supplemental Table 3 for further information and strain order. (D) Maps of Pearson correlation coefficients (*r*^2^) for pairs of Δ*gene* proteomic perturbation profiles across metabolic conditions. Strains are clustered based on respiration proteome correlations, and this strain order is held consistent across the fermentation correlations and in the addition maps in Figure S2.

A global view of these data reveal distinct strain-specific responses across all three omic planes (Figure 1C Table S3). Pairwise comparisons of the global Δ*gene* perturbation profiles reveal the overall response similarity between each strain (Figure 1D, S1A,B, and Table S4). Eight of the strains are respiratory deficient and exhibit hallmarks of the respiratory deficient response (RDR) defined in the Y3K study (Table S5) (Stefely et al., 2016). This RDR can provide important insights into the global cellular response to respiratory deficiency, but can also be normalized out in order to help reveal functionally unique phenotypes for each respiratory deficient strain. As expected, normalization of the RDR response decreased the inter-strain correlations, revealing that each gene deletion causes a largely unique global cellular response (Figure S1C). Of note, the global knockout profiles of the highly related inner membrane proteases Imp1p and Imp2p are the most highly correlated across all three omes (Figure 1D,S1A-D). Similarly, knockout of the Afg3p and Yta12p—the two subunits of the m-AAA (matrix-ATPases associated with diverse cellular activities)protease—yielded highly correlated profiles, providing an important validation of the accuracy of our mutli-omic analyses (Figure 1D,S1A-C).

We next performed outlier analyses across our dataset to systematically identify molecular alterations unique to an individual Δ*gene* strain. This analysis, which can identify functional relationships between mitoproteases and specific mitochondrial processes—and can potentially identify direct substrates —yielded 1372 Δ*gene*-specific phenotypes (Table S6). For example, the iron sulfur cluster (ISC) biogenesis proteins Isu1p and Isa1p are two of the six most highly upregulated proteins in the Δ*pim1* strain (Figure 2A), and the Δ*pim1* strain is an outlier amongst all strains for abundance increases in each of these proteins (Figure 2B,C). Isu1p is an established substrate of Pim1p (Ciesielski et al., 2016; Song et al., 2012), but, to our knowledge, Isa1p has not been associated with Pim1p. The Δ*pim1* strain also demonstrated an increase in multiple cardiolipin (CL) species (Figure S2A) suggesting that it may be a negative regulator of CL biosynthesis. In the Δ*prd1* strain, many metal ion transporters are uniquely increased during respiratory conditions (Figure S2B) leading to the hypothesis that Prd1p plays a role in regulating metal ion homoeostasis. Interestingly, the Δ*yme1* strain, which lacks the i-AAA protease complex, is an outlier for increases in Cmc1p and Coa1p—two assembly factors for complex IV (cytochrome oxidase; COX) (Figure S2C-E). The Yme1p catalytic domain faces into the mitochondrial intermembrane space (IMS) (Figure 1A) where both Cmc1p and Coa1p reside. Relatedly, the Δ*yme1* strain was an outlier for elevated levels of Cox5bp, a core COX subunit that is normally repressed during aerobic growth (Figure S2F). Collectively, these data suggest unappreciated connections between Yme1p and COX assembly and/or activity. Mpc3p (Fmp43p), a subunit of the mitochondrial pyruvate carrier (MPC), is also elevated in the Δ*yme1* strain under respiratory conditions. Mpc3p typically replaces Mpc2p as the second subunit of the MPC when yeast switch to respiratory growth from fermentation (Bender et al., 2015).These data suggest that Yme1p may help regulate mitochondrial pyruvate uptake under these growth conditions (Figure S2G). Under fermentative growth, Mgr2p—a subunit of the translocase of the inner mitochondrial membrane (TIM)complex—was uniquely increased in the Δ*imp1* and Δ*imp2* strains (Figure S2H). This corroborates a recent report suggesting that Imp1p cleaves the C-terminus of Mgr2p (Ieva et al., 2013). As a final example, our metabolomics data reveal a marked decrease in ∂-amino levulinate in the Δ*mcx1* strain (Figure S2I), corroborating a recent report that mitochondrial ClpX activates amino levulinate synthase for heme biosynthesis and erythropoiesis (Kardon et al., 2015). Many other mitoprotease functional connections can be explored via our open access data analysis and visualization tool (http://mitoproteaseprofiling.org).

**Figure 2.**
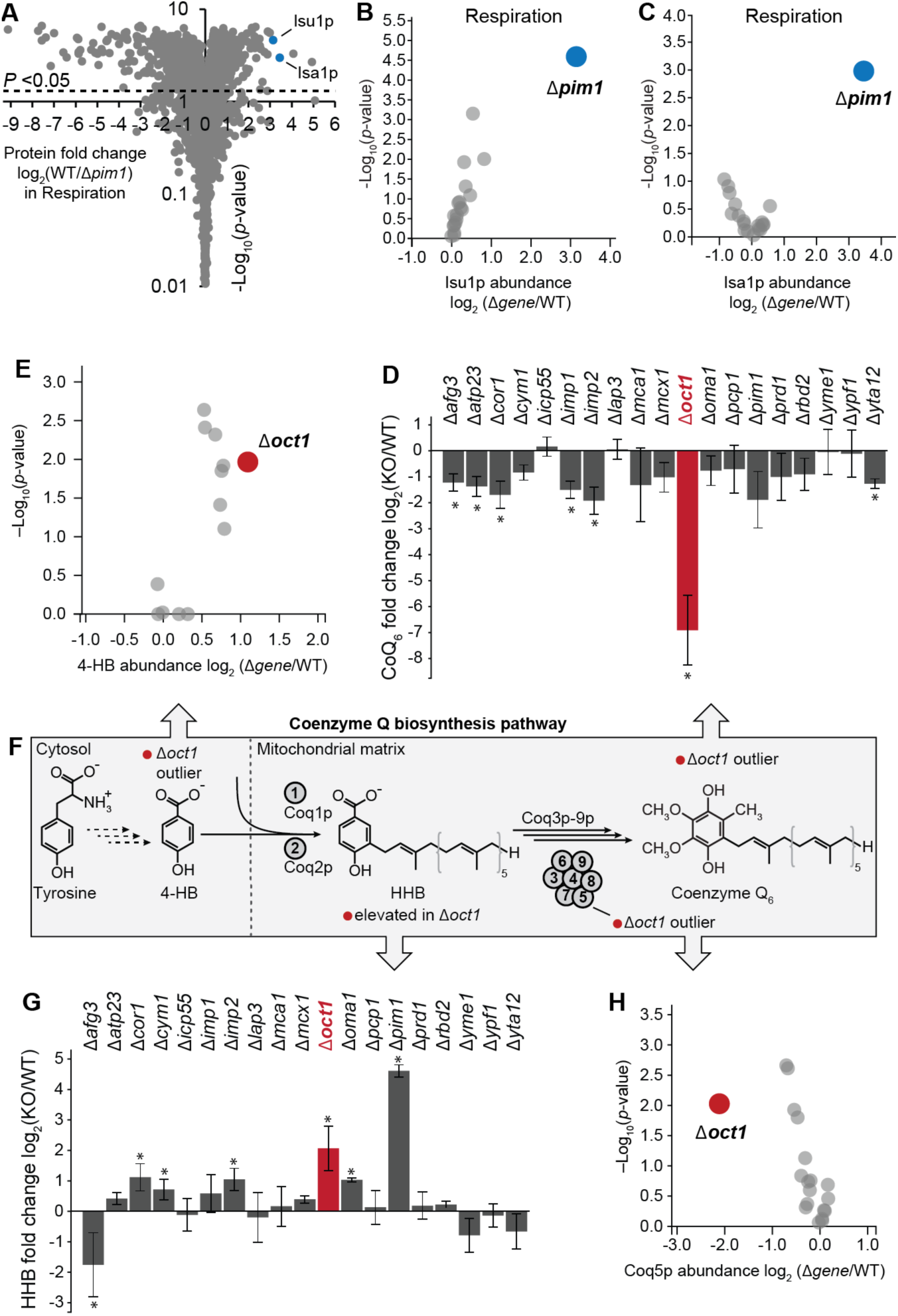
Mitoprotease profiling reveals novel connections between proteases and diverse biological processes. (A) Volcano plot showing log_2_ fold change of WT versus Δ*pim1* yeast (log_2_[Δ*pim1*/WT]) on the X-axis and significance of change (*P*-value) on the Y-axis. *P*-value was calculated with a 2-sample Student’s *t*-test. (B) Outlier analysis examining all strains for specific perturbations in Isu1p across Δ*gene* strains. Each point indicates Isu1p perturbations in a Δ*gene* strain versus WT. *P*-value was calculated with a 2-sample Student’s *t*-test. (C) Same as B for Isa1p across Δ*gene* strains (D) Bar graph showing levels of CoQ6 across Δ*gene* strains. Error bars indicate ± 1 standard deviation. * indicates a 2-sample Student’s *t*-test *P*-value less than 0.05. (E) Same as B for 4-hydroxybenzoate (4-HB) across Δ*gene* strains. (F) Abbreviated illustration of the *S. cerevisiae* CoQ6 biosynthetic pathway. Arrows pointing to other panels indicate specific measurements of individual proteins, metabolites, and lipids in this pathway. (G) Same as D for 3-hexaprenyl-4-hydroxybenzoate (HHB) levels across Δ*gene* strains. (H) Same as B for Coq5p protein abundance across Δ*gene* strains.

Our data predicted a particularly strong association between Oct1p and coenzyme Q (CoQ) biosynthesis. The Δ*oct1* strain was a clear outlier in the lipidomics data for CoQ abundance, with a >100-fold decrease under respiratory conditions (Figure 2D). Correspondingly, the Δ*oct1* strain was an outlier in the metabolomics data for an increase in 4-hydroxybenzoate (4-HB), the soluble cytosolic CoQ precursor (Figure 2E), and had elevated levels of the 3-hexaprenyl-4-hydroxybenzoate (HHB), which is formed in the mitochondrial matrix upon 4-HB import (Figure 2G) (Ashby et al., 1992). The loss of CoQ combined with the accumulation of CoQ precursors suggests that the Δ*oct1* CoQ deficiency stems from a defect in the mitochondrial matrix where Oct1p resides. The proteomics data revealed that most known Oct1p substrates were decreased in the Δ*oct1* strain (Figure S2J), consistent with their decreased stability when unprocessed, suggesting that the proteomics data could reveal candidate CoQ-related substrates that were likewise decreased in the Δ*oct1* strain. Three CoQ-related proteins fit this pattern (Coq2p, Coq5p, and Coq9p) (Figure S2K), with Coq5p being a Δ*oct1* outlier (Figure 2H). However, not all CoQ-related proteins were measured in this study (e.g., Coq10p). Moreover, Coq3p-Coq9p exist in a conserved biosynthetic complex (complex Q), and multiple studies have shown that disruption of one complex Q member can alter the abundance of others (Xie et al., 2012). As such, it is imperative to test the ability of Oct1p to process CoQ-related proteins through more direct *in vitro* and *in vivo* biochemical analyses.

### Coq5p is a direct Oct1p substrate

To test whether Oct1p is necessary for proper processing of the N-termini of CoQ proteins, we individually overexpressed Coq1p-Coq10p with C-terminal FLAG-tags in wild type (WT) and Δ*oct1* yeast. Our expectation was that substrates of Oct1p would retain eight extra residues on their N-termini when purified from the Δ*oct1* strain. We then performed α-FLAG immunoprecipitations to purify the CoQ proteins from both backgrounds, and separated the eluates by SDS-PAGE (Figure 3A). No clear size shifts were observed; however, the small size difference caused by a loss of eight amino acids might not be evident by this analysis.

**Figure 3.**
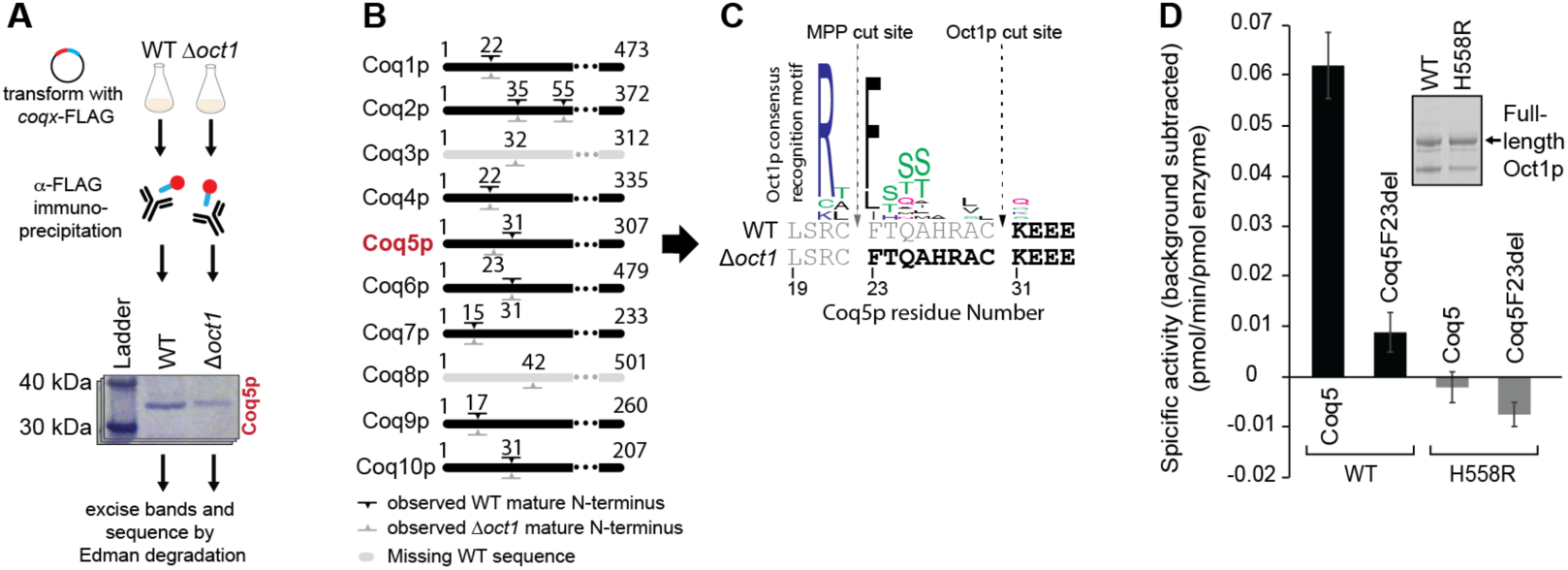
Coq5p is an Oct1p substrate. (A) Overview of process for preparing samples for Edman degradation. See Star Methods for more details. (B) Overview of N-termini observed for Coq1-10p for experiments in A. Marks indicate the N-terminus observed from proteins immunoprecipitated from WT (black) and Δ*oct1* yeast (gray). Only the N-terminus of Coq5p differed between strains. See Table S7 for all Edman results. (C) Sequence level view of Edman results for Coq5p. Residues shown in grey were not observed in the indicated strain. Sequence logo is derived from the list of known standard deviation of eigOct1p substrates described in Vögtle et al. 2011. (D) Activity of WT or catalytically dead (H558R) Oct1p against FRET peptides (see Star Methods for data acquisition and processing). Error bars indicate ± 1 standard deviation of eight replicates.

To analyze these proteins directly, we transferred the protein to a membrane, excised the protein bands and sequenced their N-termini using Edman degradation (Figure 3A,S3A-E, Table S7). For all proteins except Coq5p, the N-termini were indistinguishable when immunoprecipitated from each strain. However, the N-terminus of Coq5p from the Δ*oct1* strain was extended by exactly eight residues (Figure 3B). Moreover, these residues, along with an arginine two residues upstream, comprise a consensus Oct1p recognition motif established previously from the 14 known Oct1p substrates (Figure 3C) (Mossmann et al., 2012). These results suggest that Oct1p is responsible for processing the N-terminus of Coq5p *in vivo* following an initial processing by the mitochondrial processing protease (MPP).

To test whether Oct1p is capable of directly processing this Coq5p sequence *in vitro*, we expressed and purified full-length recombinant WT and mutant (H558R) Oct1p. The H558R mutant disrupts the Oct1p metal binding site and is ineffective at rescuing the growth deficiency of Δ*oct1* yeast (Chew et al., 1996). We incubated these constructs with Förster resonance energy transfer (FRET)-active peptides possessing C-terminal fluorophores and N-terminal quenchers, as used previously to measure the activity of the human ortholog of Oct1p, MIPEP (Marcondes et al., 2010).We tested peptides consisting of either the WT Coq5p Oct1p sequence, or a mutant version lacking the consensus phenylalanine of the Oct1p recognition sequence (F23del) (Figure 3C,S3F). WT Oct1p cleaved the WT Coq5p sequence effectively, but exhibited markedly decreased activity against Coq5p F23del (Figure 3D). The Oct1p H558R mutant was unable to cleave either peptide, demonstrating that the proteolytic activity is likely not due to a contaminating protein from the Oct1p purifications (Figure 3D). From these data, and the Oct1p outlier analyses above,we conclude that the CoQ deficiency in Δ*oct1* yeast is due to improper processing of Coq5p.

### Oct1 p processing of Coq5p supports respiratory growth and CoQ biosynthesis

To test the importance of Coq5p processing by Oct1p *in vivo*, we generated a Coq5p expression construct by mutating the same consensus motif phenylalanine as in the *in vitro* assays above (F23A).We then transformed Δ*coq5* yeast with FLAG-tagged overexpression constructs for WT Coq5p or Coq5p F23A, or an empty vector (EV) control. Both WT and F23A properly localized to mitochondria, as assessed by confocal microscopy with mitochondrial citrate synthase (Cit1p) as an endogenous control (Figure 4A). To ensure that the F23A mutant disrupted endogenous Coq5p processing, we immunoprecipitated WT and Coq5p F23A, separated the eluates by SDS-PAGE, and prepared the protein for Edman degradation sequencing as described above (Figure 4B,S4A). The F23A mutant was found predominantly in the unprocessed form (*i.e.*, eight residues longer than WT), with a lesser amount in the fully processed form (Figure 4SC, Table S7).This indicates that, similar to our *in vitro* data (Figure 3D), Oct1p likely still possesses low-level activity against the F23A mutant. Notably, this largely unprocessed F23A mutant expressed at a similar level to that of WT Coq5p in a Δ*oct1* background, which likewise cannot process Coq5p (Figure 4B). The Coq5p F23A immunoprecipitation also included a minor upper band that may represent full-length Coq5p that did not properly localize to mitochondria and/or was not processed by MPP.

**Figure 3.**
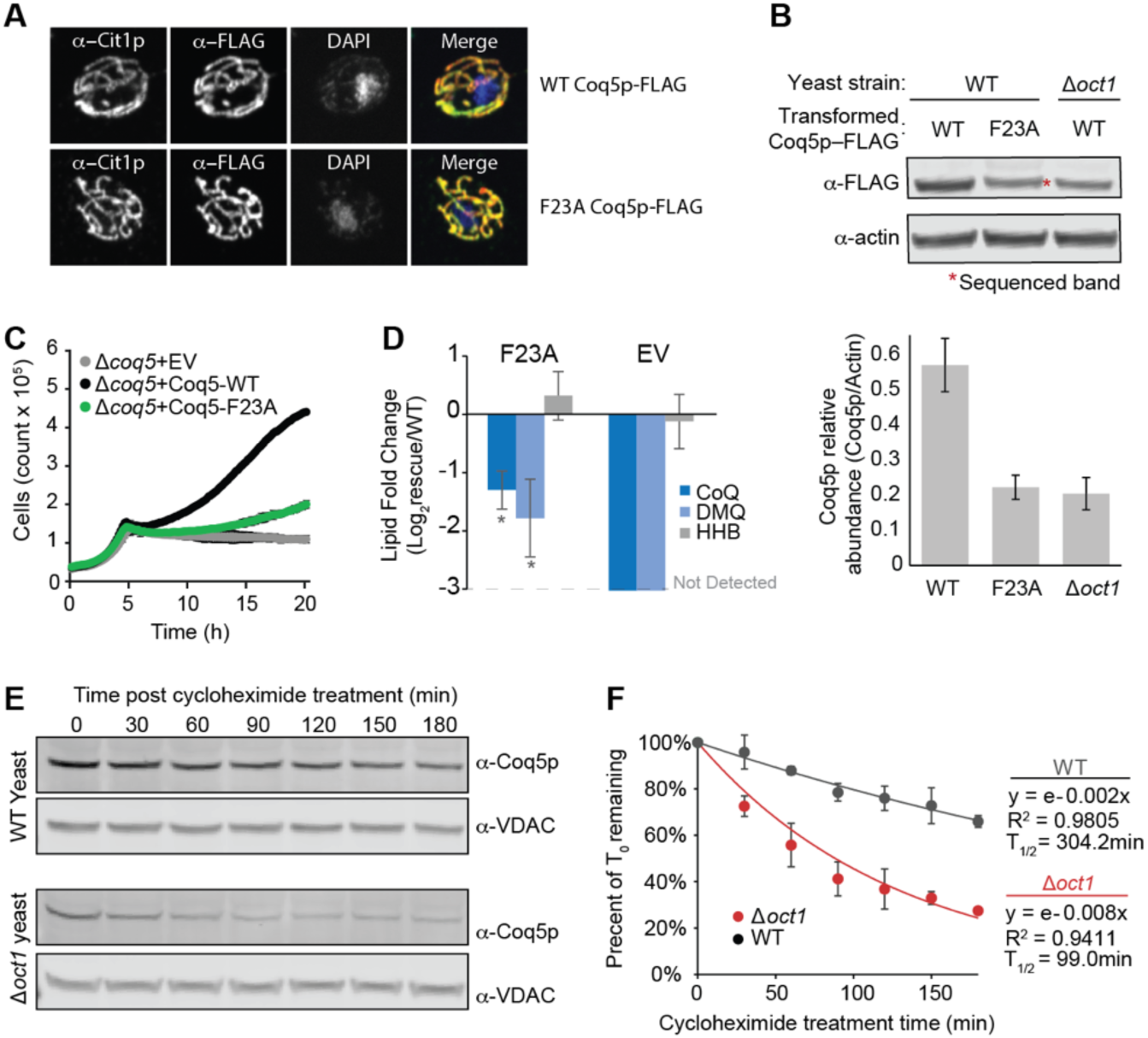
Oct1p processing stabilizes Coq5p and is essential for enabling proper CoQ_6_ production. (A) Confocal images of WT and F23A Coq5p-FLAG (red in merge) compared with citrate synthase (Cit1p) localization (green in merge) and DNA staining with DAPI (blue in merge). (B) Quantitative western blot of WT Coq5p-FLAG expressed in WT yeast (lane 1), F23A Coq5pFLAG expressed in WT yeast (lane 2), and WT Coq5p-FLAG expressed in Δ*oct1*yeast (lane 3). Lower bar graph shows relative abundance of Coq5p versus actin quantified across biological triplicate. Error bars indicate ± 1 standard deviation. (C) Growth curves of Δ*coq5* yeast rescued with indicated vector construct. (D) Targeted lipidomics measurements of CoQ-related lipids.Log_2_ fold change was calculated as a proportion of the WT rescue versus the mutant or empty vector rescue as shown in C. See methods for calculation details. Error bars indicate ± 1 standard deviation for three biological replicates. (E) Representative Western blot measuring endogenous Coq5p versus a loading control (VDAC) over time post-cycloheximide treatment. (F) Quantification of Western blots shown in E. Amount remaining was calculated as a ratio of Coq5p to VDAC normalized to T_0_ (see methods). Error bars indicate ± 1 standard deviation of four biological replicates.

We next assessed the ramifications of this improper Coq5p processing. Δ*coq5* yeast grow similarly to WT yeast under fermentation conditions, but they are respiratory incompetent, and thus cannot grow once they have exhausted their non-fermentable carbon sources (Figure 4C). Transformation of Δ*coq5* yeast with WT *coq5* rescued respiratory growth (*i.e.*, the yeast continued to past the diauxic shift when their fermentable carbon source (glucose) was exhausted). Transformation of Δ*coq5* yeast with the F23A mutant only marginally rescued this respiratory growth, consistent with the low level of properly processed Coq5p in this strain. When expressed in a WT background, Coq5p F23A had no appreciable effect on cell growth (Figure S4C). To determine whether the decreased respiratory growth was associated with altered CoQ biosynthesis, we analyzed lipid extracts from each rescue strain using LC-MS/MS. The F23A strain exhibited marked depletion of both CoQ and the direct product of Coq5p, demethoxy-CoQ (DMQ) (Figure 4D,S4D). Notably, the F23A mutant showed no difference in the early CoQ precursor, 3-hexaprenyl 4-hydroxybenzoate (HHB), whose formation does not depend on Coq5p function (Figure 4D,S4D). These results demonstrate that a single point mutation to the Oct1p recognition motif of Coq5p is sufficient to disrupt Coq5p processing, leading to CoQ deficiency and respiratory dysfunction.

Previous work has demonstrated that Oct1p processing often increases the half-lives of its substrates, perhaps by establishing a new N-terminus that is more favorable according to N-end rules (Vögtle et al., 2011). To test whether Coq5p is more stable when it is processed by Oct1p, we treated WT and Δ*oct1* yeast with cycloheximide to inhibit translation from cytoplasmic ribosomes, and collected samples over time. The results revealed a marked reduction in the stability of Coq5p in the Δ*oct1* strain where it cannot be processed by Oct1p (Figure 4F,G). Given that the change in the N-terminus from the unprocessed (phenylalanine) to the processed (lysine) form of Coq5p is predicted to provide only modest added stability according to the N-end rules (Mogk et al., 2007), the decreased half-life of unprocessed Coq5p may involve other factors (see Discussion). Purified recombinant Oct1p with or without the octapeptide had negligible differences in thermal stability *in vitro* (Figure S4E), indicating that the presence of the octapeptide itself does not add intrinsic instability to Coq5p.

Overall, these data define a unique role for Oct1p in enabling CoQ biosynthesis. Through a combination of mutli-omic profiling and biochemistry approaches, we demonstrated that Oct1p processing of Coq5p is a requisite post-translational step for ensuring proper CoQ production and respiratory competency, thereby adding a notable new function to the growing list of mitoprotease activities. Furthermore, our multi-omic profiling of 20 yeast strains under contrasting growth conditions will serve as a rich resource for additional mechanistic investigations into mitoprotease functions.

## DISCUSSION

Mitoproteases are rapidly gaining recognition as key figures in nearly every aspect of mitochondrial activities. Nonetheless, the precise functions and direct substrates of these proteins are often difficult to establish. Given their pleiotropic functions, disruption of individual mitoproteases often do not yield readily discernible cellular phenotypes that would implicate them in a distinct pathway or process. Proteases also typically do not exist in robust complexes, often have membrane associations making them difficult to purify, and have substrates that may either increase or decrease in abundance in their absence.

Recently, we devised a mass spectrometry-based multi-omic strategy to elucidate the functions of poorly characterized mitochondrial proteins (MXPs) (Stefely et al., 2016). This “Y3K” approach connects MXPs to specific pathways and processes by virtue of the unique proteomic, metabolomic, and lipidomic alterations that result from their disruption. Central to this strategy is the simultaneous analysis of many strains, thus providing contrasting perturbations from which these unique signatures can be identified. We reasoned that an in-depth profiling of nearly all mitoproteases using this approach could position each within pathways where their substrates likely exist, and might potentially reveal direct substrates themselves. Indeed, these analyses yielded more than 1000 unique outlier associations—scenarios in which a given protein, lipid, or metabolite is prominently affected by disruption of a specific mitoprotease—thus generating many functional hypotheses. These include new predicted functions for mitoproteases within pathways ranging from metal ion homeostasis, solute transport, protein import, complex assembly, cardiolipin biosynthesis, among others, which require further validation by future mechanistic work. Many other connections can be readily explored using our accompanying online data visualization tool (http://mitoproteaseprofiling.org).

Among the most striking connections in our dataset was that between Oct1p and coenzyme Q (CoQ) biosynthesis. Our multi-omic data allowed us to identify Δ*oct1* outliers for the CoQ pathway across all three omic planes: 4-HB (metabolomics), CoQ (lipidomics), and Coq5p (proteomics). These observations are supported by our recent Y3K study, in the which *oct1* was among multiple genes associated with CoQ deficiency that do not encode proteins in the canonical CoQ biosynthesis pathway (Stefely et al., 2016).Collectively, these data strongly suggested that CoQ has a particularly important relationship with this protease, and provided a clear hypothesis that could be tested biochemically. Notably, this connection could not have been easily observed by analyzing an Δ*oct1* strain alone, as the observed loss of CoQ might have been attributed to the general loss of CoQ observed in respiratory deficient strains (Branda and Isaya, 1995; Stefely et al., 2016).

Our *in vitro* and *in vivo* biochemical analyses validated our multi-omic approach, demonstrating a direct enzyme-substrate relationship with Coq5p. Although CoQ was discovered 60 years ago, many aspects of its biosynthesis remain poorly characterized, and few instances of transcriptional, post-transcriptional, or post-translational regulation of its production have been reported (Stefely and Pagliarini, in press). Intriguingly, this is the second recent report specifically demonstrating Coq5p as a recipient of such regulation: we recently showed that the regulation of Coq5p abundance by the RNA binding protein Puf3p is also crucial for proper CoQ production (Lapointe et al., 2017). Similarly, our work here demonstrates that Oct1p processing is important for Coq5p stability, and ultimately CoQ levels and respiratory competence. Given that CoQ is synthesized in the mitochondrial matrix by a biosynthetic complex (complex Q) (Floyd et al., 2016; Xie et al., 2012), these regulatory processes may be essential to set the proper complex stoichiometry and to avoid proteostatic stress resulting from unincorporated complex subunits. Indeed, recent work has linked the human Oct1p ortholog MIPEP to the mitochondrial unfolded protein response (Munch and Harper, 2016).

More than a dozen other Oct1p substrates have been identified within diverse pathways in the mitochondrial matrix, including iron homeostasis (Branda et al., 1999; Chew et al., 2000), respiratory complex formation (Fu et al., 1990), and mitochondrial DNA maintenance (Branda and Isaya, 1995).Many of these substrates have been predicted computationally and/or discovered by N-terminal proteomics (Branda and Isaya, 1995; Vögtle et al., 2009). The enhanced protein stability of Coq5p imparted by Oct1p processing is similar to most—but not all (e.g., Rip1p)— known Oct1p substrates (Vögtle et al., 2011). By and large, this added stability is attributed to the generation of new N-termini by Oct1p that can be classified as more stabilizing according to N-end rules. In the case of Coq5p, Oct1p processing results in an N--terminus featuring a lysine (secondary destabilizing residue) instead of a phenylalanine (primary destabilizing residue). Notably, this classification for lysine is based on the observation that it would typically be converted to a primary destabilizing residue by enzymes in the cytosol of eukaryotes and prokaryotes (Dougan et al., 2010); however, mitochondrial homologs of the responsible enzymes for this conversion have not been identified (Vögtle et al., 2011). Thus, and N--terminal lysine may be more stabilizing in mitochondria, consistent with the marked difference in Coq5p stability between WT and Δ*oct1* yeast. It is also possible that the octapeptide is destabilizing for other reasons, and it is noteworthy that our Coq5p F23A mutant that disrupts Oct1p processing also appears to be destabilized despite its “stabilizing” N-terminal alanine.

Our analyses indicate that other Oct1p substrates may exist that have eluded previous experimental and computational analyses. Although Coq5p possesses a similar Oct1p recognition motif to other known Oct1p substrates, its prediction as an Oct1p substrate may have been confounded by its arginine (R) residue three position upstream of its mature N--terminus. This “R-3” motif is typical for an Icp55p substrate, in which MPP would cleave the presequence one residue past the R followed by Icp55p cleavage of one additional residue (Calvo et al., 2017; Vögtle et al., 2009). However, more advanced techniques that look for sequential MPP, Oct1p motifs may have more success identifying Coq5p as an Oct1p substrate (Fukasawa et al., 2015). Additionally, an Oct1p substrate was recently identified that lacks the canonical Oct1p motif (Vögtle et al., 2011), indicating that a wider range of Oct1p substrates may exist—many of which may be downregulated in our data. Overall, these observations demonstrate the value of our unbiased mitoprotease profiling approach, and suggest that our data will provide an effective resource for further exploration of mitoprotease substrates and activities.

## SUPPLEMENTAL INFORMATION

Supplemental Information includes Supplemental Experimental Procedures, four Figures, and eight Tables.

## AUTHOR CONTRIBUTIONS

M.T.V., A.G.R., and D.J.P. conceived of the project and its design. M.T.V., A.G.R., E.C.F., N.W.K., P.D.H., M.J.D., A.J., A.U., M.J.P.R., J.J.C., D.J.P., performed experiments and/or data analysis. M.C., J.J.C., and D.J.P. provided key experimental resources and/or aided in experimental design. M.T.V. and D.J.P. wrote the manuscript.

## ACKNOWLEDGMENTS

Research reported in this publication was supported by the National Institute of General Medical Sciences of the National Institutes of Health under award numbers R01GM115591 (to D.J.P.), T32GM008505 (to A.G.R.), T32GM007215 (to M.T.V.), R35GM118110 and P41GM108538 (to J.J.C). This work was further supported by a National Science Foundation Graduate Research Fellowship DGE-1256259 (to M.T.V.). The content is solely the responsibility of the authors and does not necessarily represent the official views of the National Institutes of Health.

## SUPPLEMENTAL FIGURES

**Figure S1, related to Figure 1.**
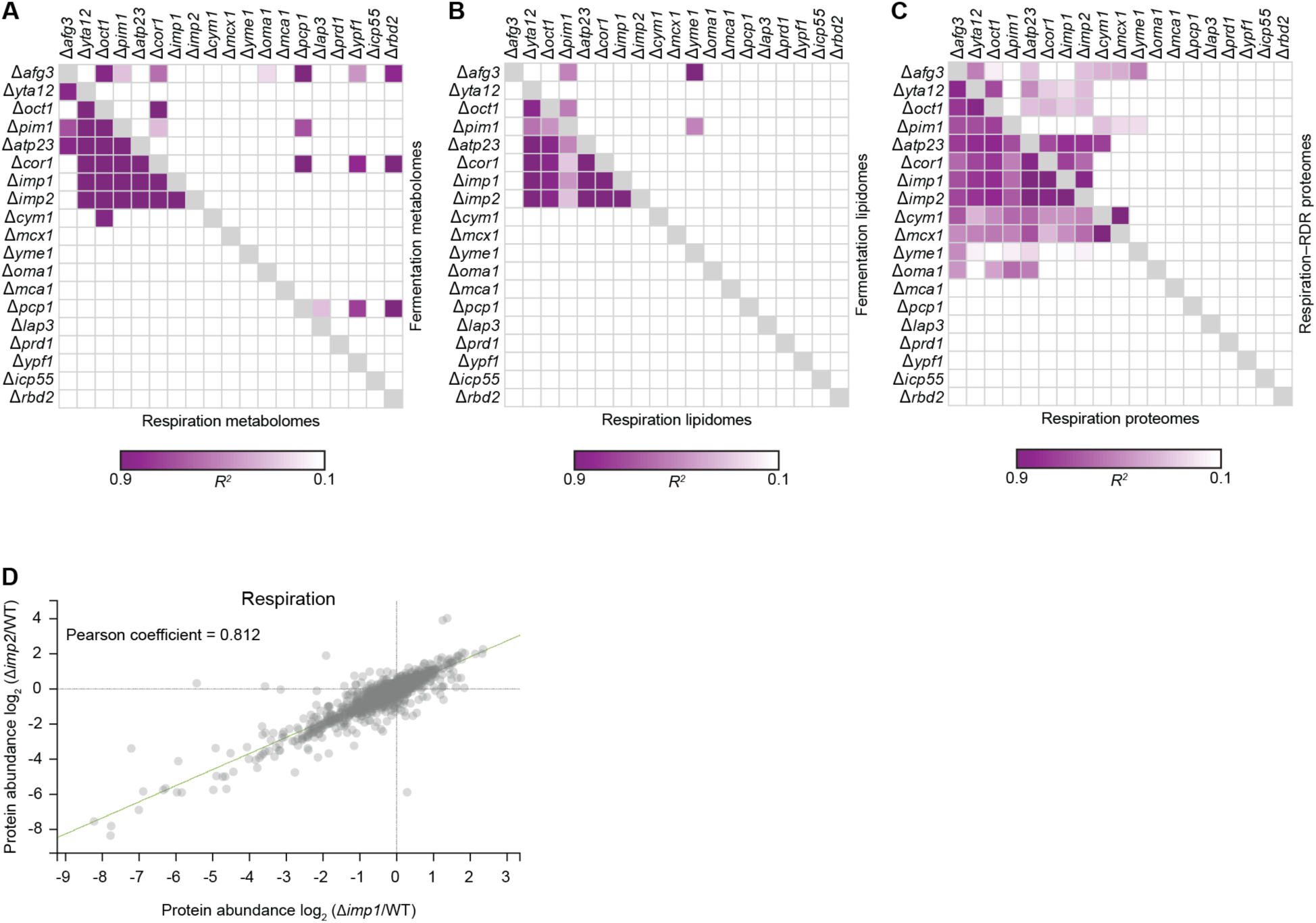
Multi-omic strain-strain correlations. (A) Strain correlation map as shown in Figure 1D for respiration and fermentation metabolite levels. (B) Strain correlation map as shown in Figure 1D for respiration and fermentation lipid levels. (C) Strain correlation map as shown in Figure 1D for respiration and respiration deficiency response (RDR)-adjusted protein levels. (D) Correlation between Δ*imp1* and Δ*imp2* proteomic signatures under respiration conditions.

**Figure S2, related to Figure 2.**
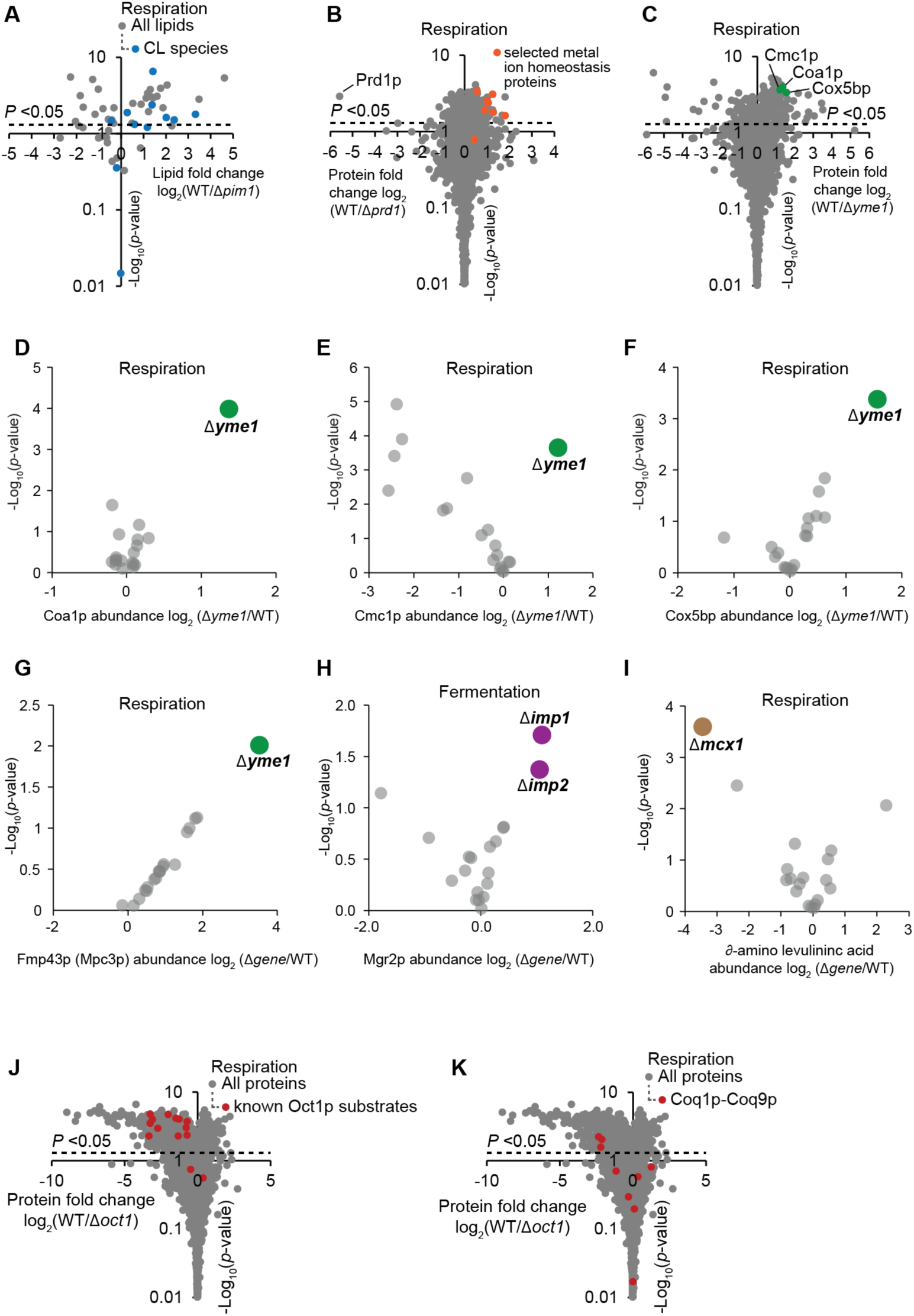
Outlier analyses reveal associations between mitoproteases and distinct mitochondrial processes. (A) Volcano plot as shown in Figure 2A for lipid species in Δ*pim1* yeast. Cardiolipin (CL) species are highlighted in blue. (B) Volcano plot as shown in Figure 2A for proteins in Δ*prd1* yeast. The following proteins related to metal ion homeostasis are highlighted in orange: Enb1p, Sit1p, Fet3p, Atx1p, Cot1p, Ftr1p, Fet5p, and Fet4p. (C) Volcano plot as shown in figure 2A for proteins in Δ*yme1* yeast. Indicated complex IV-related proteins are highlighted in green. (D) Outlier analysis as shown in Figure 2B highlighting the Coa1p abundance change in Δ*yme1* yeast. (E) Outlier analysis as shown in Figure 2B highlighting the Cmc1p abundance change in Δ*yme1* yeast. (F) Outlier analysis as shown in Figure 2B highlighting the Cox5bp abundance change in Δ*yme1* yeast. (G) Outlier analysis as shown in Figure 2B highlighting the Mcp3p abundance change in Δ*yme1* yeast. (H) Outlier analysis as shown in Figure 2B highlighting the Mgr2p abundance change in both Δ*imp1* and Δ*imp2* yeast. (I) Outlier analysis as shown in Figure 2B highlighting the ∂-amino levulinate abundance change in Δ*mcx1* yeast. (J) Volcano plot as shown in Figure 2A for proteins in Δ*oct1* yeast. Known substrates of Oct1p are highlighted in red (Vögtle et al., 2011). (K) Volcano plot as shown in Figure 2A for proteins in Δ*oct1* yeast. Proteins involved in coenzyme Q biosynthesis are highlighted in red.

**Figure S3, related to Figure 3.**
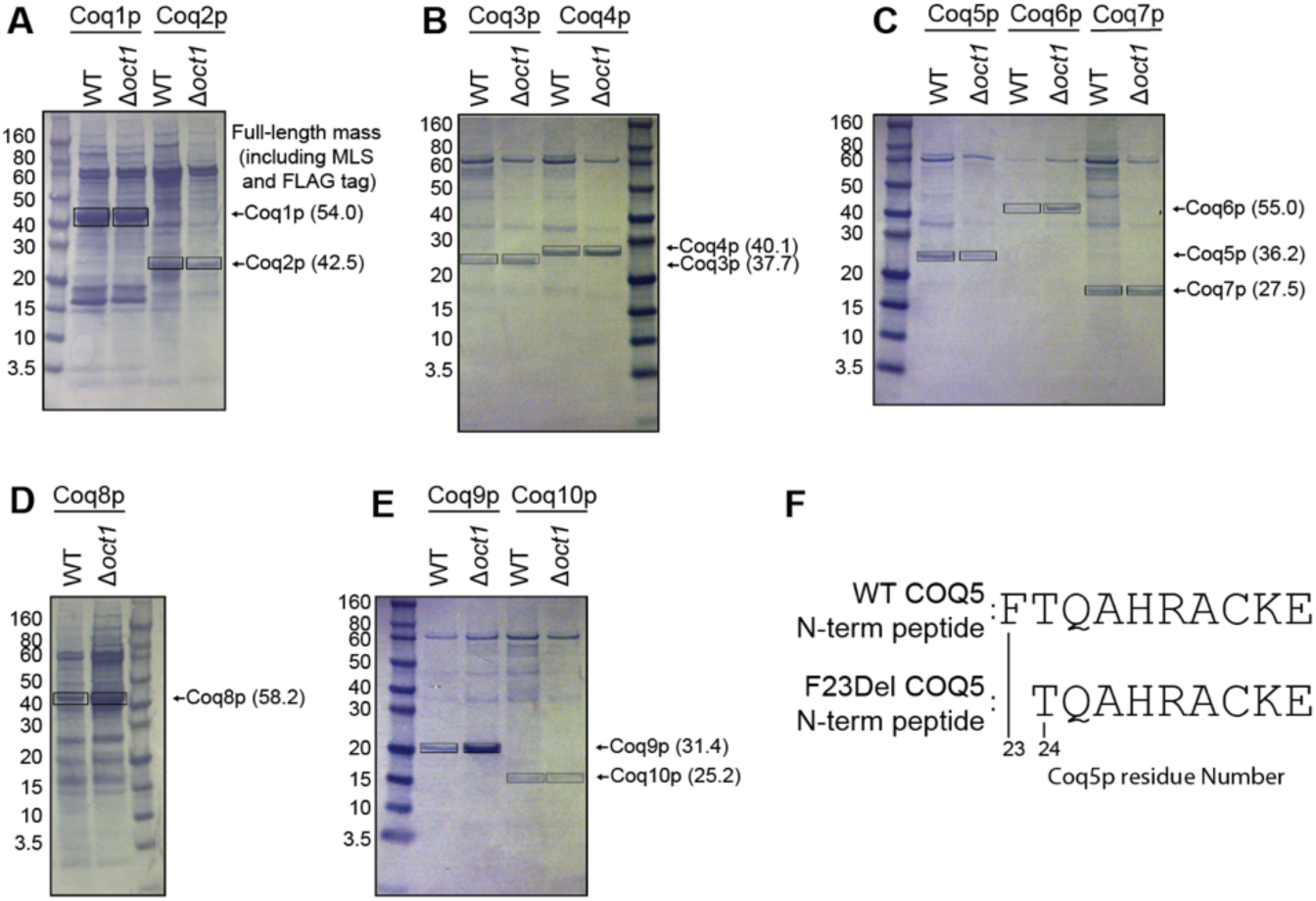
*In vitro* analyses supporting Coq5p as a substrate of Oct1p. (A) Membrane with protein transferred from SDS-PAGE analyses of Coq1p-FLAG and Coq2p-FLAG immunoprecipitations from WT and Δ*oct1* yeast. Boxes indicate bands excised for Edman sequencing. Expected full length mass of each protein including mitochondrial localization sequence (MLS) and FLAG tag are indicated in parentheses. (B) Same as A for Coq3p and Coq4p. (C) Same as A for Coq5-7p. (D) Same as A for Coq8p. (E) Same as A for Coq9p and Coq10p. (F) Sequences of the peptides used in Figure 3D.

**Figure S4, related to Figure 4.**
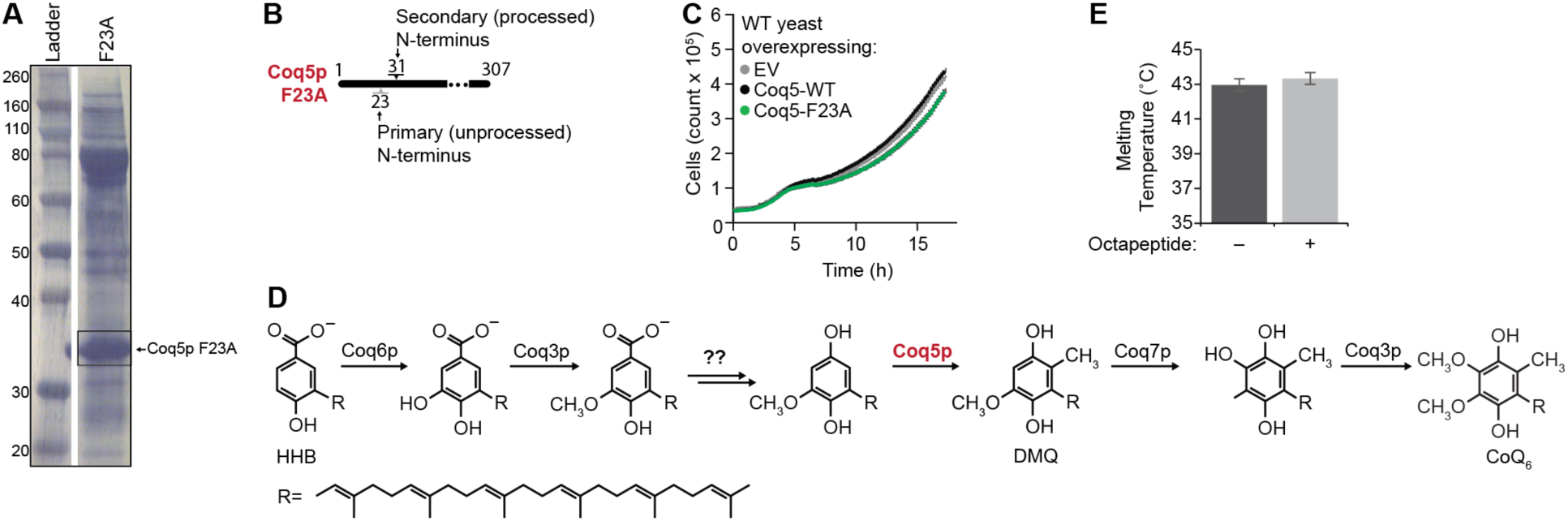
Oct1p processing stabilizes Coq5p and is essential for enabling proper CoQ6 production. (A) Coomassie-stained membrane as shown in Figure S3A-E for Coq5p F23A. Box indicates band excised for Edman sequencing. (B) Illustration as shown in Figure 3B of Edman sequencing results for Coq5p F23A. (C) Growth curve of WT yeast transformed with the same Coq5p constructs shown in Figure 4C. (D) Partial coenzyme Q biosynthesis pathway with select molecules highlighted (HHB, 3-hexaprenyl-4-hydroxybenzoate; DMQ, dimethoxy-CoQ). (E) Melting temperatures of Coq5p with and without the octapeptide (purified from *E. coli*) as measured by differential scanning fluorimetry (DSF) (see Star Methods for further information). Error bars indicate ± 1 standard deviation.

## SUPPLEMENTAL TABLE LEGENDS

**Supplementary Table 1. Knockout yeast strains.**

Table of single-gene deletion (Δ*gene*) yeaststrain s investigated in this study and their harvest culture densities. For each gene deleted in a strain studied. The second tab shows the culture densities upon harvest (growth phenotypes), and it includes the systematic yeast gene name, the standard gene name, predicted human homologs, the culture densities at time of harvest, the average of these densities (mean, n = 3), and the corresponding standard deviations.

**Supplementary Table 2. Profiled biomolecules.**

Table of all molecules profiled in the study. Includes molecule type (protein, lipid, or metabolite), molecule name, standard gene name (for proteins) or standard lipid name, systematic gene name (for proteins), and UniProt ID (for proteins). For lipids, the numbers in parentheses indicate the number of carbons in the acyl tail(s) and the number of carbon-carbon double bonds in the chains (carbons:double_bonds).

**Supplementary Table 3. Quantitative dataset.**

Table containing quantitative measurements and descriptive statistics used throughout the study. Average fold changes in molecule abundances (mean log_2_[Δ*gene*/WT], n = 3) for all strains and all molecules in the respiration and fermentation datasets are shown on tabs labeled ‘KO vs WT_Resp (ΔLFQ)’ and ‘KO vs WT_Ferm (ΔLFQ)’. Corresponding standard deviations, and *P*-values (2-tailed t-test [homostatic]) for all measured fold changes are shown on separate tabs labeled accordingly with ‘(Std. Dev.)’ and ‘(*P*-Value).’ Each tab contains a table with rows corresponding to molecules and columns corresponding to the single gene knockout (Δ*gene*) strains profiled in the study.

**Supplementary Table 4. Δgene–Δgene perturbation profile correlations.**

Table of Δ*gene*–Δ*gene* perturbation profile correlations (Pearson coefficients). Includes the gene knocked out of ‘strain one’ in the pairwise comparison, the gene knocked out of ‘strain two’ in the pairwise comparison, the Pearson coefficient, and the ‘Ome’ (proteome, lipidome, or metabolome) used for the regression analysis. Coefficients are only reported for Δ*gene*–Δ*gene* pairs meeting the criteria outlined in the Methods under the heading ‘Regression analysis of phenotype changes’. Separate tabs are included for the respiration (resp), fermentation (ferm), and RDR-adjusted respiration (resp-RDR) datasets.

**Supplementary Table 5. Respiration deficient strains versus wild type.**

Table of average fold change in molecule abundances (mean log_2_[RD strains/WT]). Includes molecule identifiers (including UniProt IDs, symbols, and systematic names for proteins), fraction of RD strains showing consistent perturbation of each molecule, RDR score (see Methods), and average fold change (mean log_2_[RD strains/WT]).

**Supplementary Table 6. Δgene-specific phenotypes.**

Table of unique Δ*gene*-phenotype relationships identified in this study. Includes molecule name (standard gene name followed by systematic gene name for all proteins), yeast deletion strain (standard gene name), calculated Euclidean distance, and associated growth condition (respiration or fermentation).

**Supplementary Table 7. Edman results corresponding to Figures 3 and 4.**

Table summarizing Edman results observed including information on the protein that was sequenced,in which yeast background it was expressed, which figure the results correspond to, and the expected and observed protein masses. This table also includes the full-length sequence of each protein (including a C-terminal FLAG-tag), the sequence data, and where that sequence data aligns within the full-length protein sequence.

**Supplementary Table 8. Primers table**

List of the primers and their sequences used in this study.

## STAR METHODS

### CONTACT FOR REAGENT AND RESOURCE SHARING

Further information and requests for resources and reagents should be directed to and will be fulfilled by the Lead Contact, Dave Pagliarini.

### EXPERIMENTAL MODEL AND SUBJECT DETAILS

*Escherichia coli* strain DH5α (NEB) was used for all cloning applications and grown at 37 °C in LB media with antibiotics. *Escherichia coli* strain BL21-CodonPlus (DE3)-RIPL (Agilent) was used for all protein expression and purification purposes. See methods below for more details. Yeast work was done in *Saccharomyces cerevisiae* strain BY4742 was used. See methods below for more details.

## METHOD DETAILS

### Cell culturing

The parental (WT) *Saccharomyces cerevisiae* strain for this study was the haploid MATalpha BY4742. Single gene deletion (Δ*gene*)derivatives of BY4742 were either obtained through the gene deletion consortium (Giaever et al., 2002) (Thermo #YSC1054) or made in-house using a *KanMX* deletion cassette to match those in the consortium collection. All gene deletions were confirmed by either proteomics (significant decrease in the encoded protein) or a PCR assay. Δ*gene* strains made in-house were also confirmed by gene sequencing.

Single lots of yeast extract (‘Y’) (Research Products International, RPI), peptone (‘P’) (RPI), agar (Fisher), dextrose (‘D’) (RPI), glycerol (‘G’) (RPI), and G418 (RPI) were used for all media. YP and YPG solutions were sterilized by automated autoclave. G418 and dextrose were sterilized by filtration (0.22 μm pore size, VWR) and added separately to sterile YP or YPG. YPD+G418 plates contained yeast extract (10 g/L), peptone (20 g/L), agar (15 g/L), dextrose (20 g/L), and G418 (200 mg/L). YPD media (fermentation cultures) contained yeast extract (10 g/L), peptone (20 g/L), and dextrose (20 g/L). YPGD media (respiration cultures) contained yeast extract (10 g/L), peptone (20 g/L), glycerol (30 g/L) and dextrose (1 g/L).

Yeast from a −80 °C glycerol stock were streaked onto YPD+G418 plates and incubated (30 °C, ~60 h). Starter cultures (3 mL YPD) were inoculated with an individual colony of yeast and incubated (30 °C, 230 r.p.m., 10–15 h). A WT culture was included with each set of Δ*gene* strain cultures (19 Δ*gene* cultures and 1 WT culture). Cell density was determined by optical density at 600 nm (OD_600_) as described (Hebert et al., 2013). YPD or YPGD media (100 mL media at ambient temperature in a sterile 250mL Erlenmeyer flask) was inoculated with 2.5 × 10^6^ yeast cells and incubated (30 °C, 230 r.p.m.). Samples of the YPD cultures were harvested 12 h after inoculation, a time point that corresponds to early fermentation (logarithmic) growth. Samples of YPGD cultures were harvested 25 h after inoculation, a time point that corresponds to early respiration growth.

### Liquid chromatography tandem mass spectrometry (LC–MS/MS) proteomics

1 × 10^8^ yeast cells were harvested by centrifugation (3,000 *g*, 3 min, 4 °C), the supernatant was removed, and the cell pellet was flash frozen in N_2(l)_ and stored at −80°C. Yeast pellets were resuspended in 8 M urea, 100 mM tris (pH = 8.0). Yeast cells were lysed by the addition of methanol to 90%, followed by vortexing (~30 s). Proteins were precipitated by centrifugation (12,000 *g*, 5 min). The supernatant was discarded, and the resultant protein pellet was resuspended in 8 M urea, 10 mM tris(2-carboxyethyl)phosphine (TCEP), 40 mM chloroacetamide (CAA)and 100 mM tris (pH = 8.0). Sample was diluted to 1.5 M urea with 50 mM tris and digested with trypsin (Promega) (overnight, ~22 °C) (1:50, enzyme/protein). Samples were desalted using Strata X columns (Phenomenex Strata-X Polymeric Reversed Phase, 10 mg/mL). Strata X columns were equilibrated with one column volume of 100% acetonitrile (ACN), followed by 0.2% formic acid. Acidified samples were loaded on column, followed by washing with three column volumes of 0.2% formic acid or 0.1% trifluoroacetic acid (TFA). Peptides were eluted off the column by the addition of 500 μL 40% ACN with either 0.2% formic acid or 0.1% TFA and 500 μL 80% ACN with either 0.2% formic acid or 0.1% TFA. Peptide concentration was measured using a quantitative colorimetric peptide assay (Thermo). LC–MS/MS analyses were performed using previously described methodologies (Hebert et al., 2014a; Richards et al., 2015b)

#### Data analysis

Raw data files were acquired in one batch of 60 (3 biological replicates of 19 Δgene strains and 1 WT strain) with time between LC–MS analyses minimized to reduce run-to-run variation. This batch of raw data files were subsequently processed using MaxQuant (Cox and Mann, 2008) (Version 1.5.0.25). Searches were performed against a target-decoy(Elias and Gygi, 2007) database of reviewed yeast proteins plus isoforms (UniProt, downloaded January 20, 2013) using the Andromeda (Cox et al., 2011) search algorithm. Searches were performed using a precursor search tolerance of 4.5 p.p.m. and a product mass tolerance of 0.35 Da. Specified search parameters included fixed modification for carbamidomethylation of cysteine residues and a variable modification for the oxidation of methionine and protein N-terminal acetylation, and a maximum of two missed tryptic cleavages. A 1% peptide spectrum match (PSM) false discovery rate (FDR) and a 1% protein FDR was applied according to the target-decoy method. Proteins were identified using at least one peptide (razor + unique). Proteins were quantified using MaxLFQ with a label-free quantification (LFQ) minimum ratio count of 2. LFQ intensities were calculated using the match between runs feature, and MS/MS spectra were not required for LFQ comparisons. Missing values were imputed where appropriate for proteins quantified in ≥50% of MS data files in a batch. Proteins not meeting this requirement were omitted from subsequent analyses. Imputation was performed on a replicate-by-replicate basis. For each replicate MS analysis a normal distribution with mean and s.d. equivalent to that of the lowest 1% of measured LFQ intensities was generated. Missing values were filled in with values drawn from this distribution at random.. Replicate protein LFQ values from corresponding Δ*gene* or WT strains were pooled, log_2_ transformed, and averaged (mean log_2_[strain], *n* = 3). Average Δ*gene* LFQ intensities were normalized against their appropriate WT control (mean log_2_[Δ*gene*/WT], *n* = 3) and a 2-tailed *t*-test (homostatic) was performed to obtain *P* values.

### LC–MS lipidomics

1 × 10^8^ yeast cells were harvested by centrifugation (3,000 *g*, 3 min, 4 °C), the supernatant was removed, and the cell pellet was flash frozen in N_2(1)_ stored at−80°C Frozen yeast pellets (1 × 10^8^ cells) were thawed on ice and mixed with glass beads (0.5 mm diameter, 100 μL). CHCl_3_/MeOH (1:1, v/v, 4 °C) (900 μL) was added and vortexed (2 × 30 s). HCl (1 M, 200 μL, 4 °C) was added and vortexed (2 × 30 s). The samples were centrifuged (5,000 *g*, 2 min, 4 °C) to complete phase separation. 400 μL of the organic phase was transferred to a clean tube and dried under Ar_(g)._ The organic residue was reconstituted in ACN/IPA/H_2_O (65:30:5, v/v/v) (100 μL) for LC–MS analysis.

LC–MS analysis was performed on an Ascentis Express C18 column held at 35 °C (150 mm × 2.1 mm × 2.7 μm particle size; Supelco) using an Accela LC Pump (500 μL/min flow rate; Thermo). Mobile phase A consisted of 10 mM ammonium acetate in ACN/H_2_O (70:30, v/v) containing 250 μL/L acetic acid. Mobile phase B consisted of 10 mM ammonium acetate in IPA/ACN (90:10, v/v) with the same additives. Initially, mobile phase B was held at 50% for 1.5 min and then increased to 95% over 6.5 min where it was held for 2 min. The column was then reequilibrated for 3.5 min before the next injection. 10 μL of sample were injected by an HTC PAL autosampler (Thermo). The LC system was coupled to a Q Exactive mass spectrometer (Build 2.3 SP2) by a HESI II heated ESI source kept at 325 °C (Thermo). The inlet capillary was kept at 350 °C, sheath gas was set to 35 units, and auxiliary gas to 15 units, and the spray voltage was set to 3,000 V. Several scan functions were used to achieve optimal data acquisition for different lipid classes. For phospholipids, MS^1^ (MS scan of precursor ions without fragmentation) data were acquired from 1–9 min at a resolving power of 35,000 with AGC the target set to 1 × 10 ^6^, mass range to 500–900 Th, and maximum injection time to 250 ms. For fatty acids and lyso species (lipids lacking a fatty acyl tail) MS^1^ data were acquired from 0–3 min at a resolving power of 17,500 with the AGC targetset to 5 × 10^5^, mass range to 220–600 Th, and maximum injection time to 100 ms. For cardiolipins, MS^1^ data were acquired from 6.5–9.5 min at a resolving power of 17,500 With the AGC target setto5 × 10, mass range to 1320–1500 Th, and maximum injection time to 250 ms. For cytidine diacylglycerols, MS^1^ data were acquired from 1–4.5 min at a resolving power of 17,500 with the AGC target set to 5 × 10^5^, mass range to 920–1050 Th, and maximum injection time to 250 ms. Quantitation for all of these species was performed by integrating the MS^1^ peak areas of either the [M–H]^−^ or [M+Ac]^−^ ions. Coenzyme Q_6_ and demethoxycoenzyme Q_6_ were monitored from 4.7 to 5.8 min by positive ion tandem mass spectrometry using the 591.44 → 197.08 Th and 561.43 → 167.07 Th transitions at a normalized collision energy of 27 units, a resolving power of 17,500, a maximum injection time of 250 ms, and an isolation width of 1.5 Th. For some follow-up studies MS^1^ spectra were acquired from 200–1550 *m*/*z* supplemented with scheduled targeted scan modes to quantify key CoQ intermediates in their optimal polarity.

#### Data analysis

Peaks were automatically integrated using TraceFinder software (Thermo) and all integrations were checked manually. Missing values from undetected peaks were imputed and imputation was performed on a replicate-by-replicate basis. Total measured ion current from peaks quantified within replicate MS analyses was normalized to a corresponding WT control using a two-step procedure. First, to account for differences in cardiolipin extraction efficiency, summed cardiolipin intensities were normalized to equal the summed intensity of corresponding cardiolipin species in the WT control. All other lipid intensities were then normalized to equal the summed intensity of non-cardiolipin species in the same control. Replicate lipid intensities from corresponding Δ*gene* or WT strains were pooled, log_2_ transformed, and averaged (mean log_2_[strain], *n* = 3). Mean intensities were then normalized to WT (mean log_2_[Δ*gene*/WT], *n* = 3) and a 2-tailed *t*-test (homostatic) was performed to obtain *P*-values.

### Gas chromatography–mass spectrometry (GC–MS) metabolomics

1 × 10^8^ yeast cells yeast cells were isolated by rapid vacuum filtration onto a nylon filter membrane (0.45 μm pore size, Millipore) using a Glass Microanalysis Filter Holder (Millipore), briefly washed with phosphate buffered saline (1 mL), and immediately submerged into ACN/MeOH/H_2_O (2:2:1, v/v/v, 1.5 mL, pre-cooled to -20 °C) in a plastic tube. The time from sampling yeast from the culture to submersion in cold extraction solvent was less than 30 s. Tubes with the extraction solvent, nylon filter, and yeast were stored at -80 °C before analysis.

Tubes with yeast extract (also still containing insoluble yeast material and the nylon filter) were thawed at room temperature for 45 min., vortexed (~15 s), and centrifuged at room temperature (6,400 r.p.m., 30 s) to pellet insoluble yeast material. Yeast extract (25 μL aliquot) and internal standards (25 μL aqueous mixture of isotopically labeled alanine-2,3,3,3-d_4_, adipic acid-d_10_, and xylose-^13^C_5_ acid, 5 p.p.m. in each) were aliquoted into a 2 mL plastic tube and dried by vacuum centrifuge (~1 h). The dried metabolites were resuspended in pyridine (25 μL) and vortexed. 25 μL of *N*-methyl-*N*-trimethylsilyl]trifluoroacetamide (MSTFA) with 1% tri methylch loros i lane (TMCS) was added, and the sample was vortexed and incubated (60 °C, 30 min). Samples were then transferred to glass autosampler vials and analyzed using a GC–MS instrument comprising a Trace 1310 GC coupled to a Q Exactive Orbitrap mass spectrometer. For the yeast metabolite extracts a linear temperature gradient ranging from 50 °C to 320 °C was employed spanning a total runtime of 30 min. Analytes were injected onto a 30-m TraceGOLD TG-5SILMS column (Thermo) using a 1:10 split at a temperature of 275 °C and ionized using electron ionization (EI). The mass spectrometer was operated in full scan mode using a resolution of 30,000 (*m*/Δ*m*) relative to 200 *m/z*.

#### Data analysis

The resulting GC–MS data were processed using an in-house-developed software suite (https://github.com/coongroup/Y3K-Software). Briefly, all *m/z* peaks are aggregated into distinct chromatographic profiles (i.e., feature) using a 10 p.p.m. mass tolerance. These chromatographic profiles are then grouped according to common elution apex (i.e., feature group). The collection of features (i.e., *m*/*z* peaks) sharing a common elution apex, therefore, represent an individual electron ionization (EI)–MS spectrum of a single eluting compound. The EI–MS spectra were then compared against a matrix run and a background subtraction was performed. Remaining EI–MS spectra are then searched against the NIST 12 EI–MS library and subsequently subjected to a high-resolution filtering (HRF) technique as described elsewhere. EI–MS spectra that were not identified were assigned a numeric identifier. Feature intensity, which was normalized using total metabolite signal, was used to estimate metabolite abundance. Replicate metabolite intensities from corresponding Δ*gene* or WT strains were pooled, log_2_ transformed, and averaged (mean log_2_[strain], *n* = 3). Average Δ*gene* metabolite intensities were normalized against their appropriate WT control (mean log_2_[Δ*gene*/WT], *n* = 3) and a 2-tailed *t*-test was performed to obtain *P*-values.

### Δ*gene*-specific phenotype detection

For each profiled molecule (in both respiration and fermentation growth conditions) we separated potential Δ*gene*-specific measurements into two groups: positive log_2_ fold change (log_2_[Δ*gene*/WT]) and negative log_2_ fold change. These two sets were then plotted individually with log_2_ fold change and −log_10_(*P*-value[two-sided Student’s *t*-test]) along the *x* and *y* axes, respectively. Data were normalized such that the largest log_2_ fold change and largest −log_10_(*P*-value) were set equal to 1. Considering the three largest fold changes where *P* < 0.05, we calculated the Euclidean distance to all neighboring data points and stored the smallest result. A requirement was imposed that all considered ‘neighbors’ have a smaller fold change than the data point being considered. It is anticipated that data points corresponding to Δ*gene*-specific phenotypes will be outliers in the described plots and have large associated nearest-neighbor Euclidean distances. The described routine yielded three separate distances, the largest of which was stored for further analysis. We set a cutoff for classification as a ‘Δ*gene*-specific phenotype’ at a Euclidean distance of 0.70.

### Regression analysis of phenotype changes

#### Regression analysis of Δgene–Δgene perturbation profiles

For all pairwise combinations of Δ*gene* strains from the same growth condition linear regression analysis was conducted on protein, lipid, and metabolite perturbation profiles, respectively. Fold change (FC) measurements (mean log_2_[Δ*gene*/WT], *n* = 3) from molecules where FC > 0.7 and *P* < 0.05 were used and a minimum of 20 proteins, 10 metabolites, and 5 lipids, respectively, were required. These measurements were fit to a line and the associated Pearson correlation coefficient was reported. Coefficients carrying negative signs were set to 0. For pairs of Δ*gene* strains lacking a sufficient number of molecules that met the aforementioned criteria, the Pearson coefficient was reported as 0. Hierarchical clustering of Δ*gene*–Δ*gene* correlations was performed as described below. *Respiration deficiency response (RDR) abundance adjustment.* All Δ*gene* strains grown under respiration conditions were classified as respiration deficient (RD) (8) or respiration competent (RC) (111), based on observation of a common perturbation profile signature. For all molecules profiled within RD Δ*gene* strains an RDR score was calculated. This metric represents the proportion of RD Δ*gene* replicates over which the molecule was consistently perturbed, relative to all RD Δ*gene* replicates where the molecule was quantified. Considering all RD Δ*gene* strains, 982 molecules produced an RDR score > 0.95 (consistently perturbed across more than 95% of RD Δ*gene* replicates where quantified) and were subsequently classified as RDR-associated. For each RDR-associated molecule, individual RD Δ*gene* strain measurements were mean normalized and stored. These RDR-adjusted measurements were then used in described respiration–RDR analyses.

#### Regression analysis of RDR-adjusted Δgene–Δgene perturbation profiles

For all RD Δ*gene* strains linear regression analysis was performed pairwise on RDR-adjusted protein perturbation profiles. Fold change measurements from molecules where FC > 0.7 and *P* < 0.05 (*P*-value before RDR adjustment) were used and a minimum of 20 proteins was required. Correlations and clustering were otherwise conducted as described above.

### Hierarchical clustering

All hierarchical clustering performed in this study was done in Perseus. For all clustering operations Spearman correlation was used with average linkage, preprocessing with *k*-means, and the number of desired clusters set to 300 for both rows and columns. For clustering of Δ*gene* perturbation profiles, clustering was performed separately for fermentation and respiration data sets, and column-wise cluster order for fermentation and respiration data sets was generated using only protein fold change profiles. Column ordering was then applied to metabolite and lipid fold change data sets from the corresponding growth condition and row-wise clustering was conducted. GO term enrichment was performed in Perseus. *P* values were obtained from a Fisher’s exact test, adjusted for multiple hypothesis testing^39^ and reported where *P* < 0.05.

For the analysis of Δ*gene*–Δ*gene* correlations, clustering was performed on respiration protein perturbation profile correlation data, and the resultant ordering was applied to Δ*gene*–Δ*gene* correlation data sets from all other omes and growth conditions for parallel visual display. The same clustering process was carried out for the analysis of Δ*gene*–Δ*gene* correlations of RD Δ*gene* strains following RDR adjustment.

### Cloning for Edman sequencing

Coq1-10 was cloned into p416 GPD with a C-terminalFLAG-tagusing SpeI as the forward restriction site and MluI as the reverse (Mumberg et al., 1995) (see primers table (Table S8) for primers used for cloning). Vector digestion was done at 37 C for 1 hour followed by CIP treatment. Insert amplification was done using Accuprime Pfu polymerase (Invitrogen, USA). Insert and vector were ligated and transformed into DH5α *Escherichia coli*. Plasmid minipreps were performed and recombinants were confirmed by sequencing.

### Edman sequencing

Vectors harboring overexpression constructs of Coq1-10 were transformed into either WT (BY4742 see key resources table) or Δ*oct1*(BY4742 Δ*oct1* see key resources) yeast. Single colony transformants were picked into 5 mL of overnight –URA dropout media containing 2% glucose. The following day, 2.5 × 10^7^ yeast cells were inoculated into 1 L of –URA drop out media containing 0.1% glucose and 3% glycerol. This was allowed to grow for 25 hours at 30°C with 220 RPM shaking.

The following day, yeast spheroplasts were prepared as previously described (Boldogh and Pon, 2007). In brief, yeast were pelleted at 4000xG for 5 min at 25°C in 1 L bottles. Yeast were washed once in Milli-Q water while being transferred to a 50 mL falcon tube. Yeast were again pelleted at 4000xG for 5 min at 25°C. Supernatant was discarded and protein pellet wet weight was weighed. Yeast pellet was resuspended in 25 mL of pretreatment buffer (0.1-M Tris–SO_4_, pH 9.4, 10-mM DTT). This was incubated at 30°C with 220 RPM shaking for 15 min before pelleting down and discarding the supernatant. From here, yeast were resuspended in SP buffer (1.2 M Sorbitol, 20 mM KPi, pH 7.4). 100 μL of 100 mg/mL zymolyase (see Key Resources) was added per gram of yeast pellet measured previously. This was left to incubate at 30°C with 220 RPM shaking for 40 min. Afterwards, the yeast was pelleted at 4500xG for 5 min at 4°C before washing and re-pelletting the spheroplasts with fresh SP buffer with protease inhibitors. Spheroplasts were resuspended in 30 mL of lysed in lysis buffer with protease inhibitors (10 mM Tris-Cl pH=8.0, 1 mM EDTA, 0.5 mM EGTA, 140 mM NaCl, 1% Triton X-100, 0.1% sodium deoxycholate, and 0.1% Sodium dodecyl sulfate) and sonicated for a total of 1 min at 75% amplitude with a 1/4 inch tip in 15 second bursts. Lysate was clarified by centrifugation at 15,000 x G for 30 min at 4°C

Clarified lysate was added to 300 μL FLAG slurry (see Key Resources) after slurry had been equilibrated with wash buffer (10 mM Tris-Cl pH=8.0, 1 mM EDTA, 0.5 mM EGTA, 140 mM NaCl, 0.1% Triton X-100, 0.01% sodium deoxycholate, and 0.01% Sodium dodecyl sulfate). Clarified lysate slurry mix was placed on an end-over-end shaker for 1 hour at 4°C before slurry was pelleted at 1000 xG for 3 min at 4°C.

Pelleted slurry was resuspended in wash buffer and transferred to a 1.5 mL microcentrifuge tube. Slurry was washed 3 times with wash buffer before protein was eluted into 300 μL of elution buffer (10 mM Tris-Cl pH=8.0, 1 mM EDTA, 0.5 mM EGTA, 140 mM NaCl, 1% Triton X-100, 0.1% sodium deoxycholate, 0.1% Sodium dodecyl sulfate, and 0.2 mg/mL FLAG peptide).

30 μL of elution was run on a gel and transferred to an Immobilon-P^SQ^ membrane. Membrane was stained with Coomassie and the proper band was excised. Excised band was sent for Edman analysis at the University of Iowa Protein Sequencing Facility.

### Protein prep for Oct1p

PIPE cloning was used to generate pVP68K vectors encoding full length Oct1p and a version of Oct1 p with the H558R mutation previously shown to disrupt Oct1p’s processing ability (Chew et al., 1996). These vectors contained an 8x C-terminal His-tag followed by MBP and a TEV cleavage site. These constructs were expressed in 3 L of *E. coli* (BL21[DE3]-RIPL strain) by 1 mM IPTG overnight at 16°C. Cells were isolated and resuspended in 100 mL of lysis buffer (50 mM HEPES, 200 mM NaCl, 10% glycerol, 5 mM BME, 0.25 mM PMSF, 1 mg/mL lysozyme (Sigma), pH 8.0). Cells were lysed by sonication (4 °C, 2 × 20 s), and the lysate was clarified by centrifugation (15,000 *g*, 30 min, 4 °C). The clarified lysate was mixed with 5 mL bed volume cobalt IMAC resin (Talon resin) and incubated (4 °C, 1 h). The resin was pelleted by centrifugation (700 *g*, 2 min, 4 °C) and washed three times (~10 resin bed volumes each) with wash buffer (50 mM HEPES, 200 mM NaCl, 10% glycerol, 5 mM BME, 0.25 mM PMSF, 10 mM imidazole, pH 8.0). His-tagged protein was eluted with elution buffer (50 mM HEPES, 200 mM NaCl, 10% glycerol, 5 mM BME, 0.25 mM PMSF, 100 mM imidazole, pH 8.0). The eluted protein was concentrated with a 50-kDa MW-cutoff spin filter (Merck Millipore Ltd.) and exchanged into storage buffer (50 mM HEPES, 200 mM NaCl, 10% glycerol, 5 mM BME, 0.25 mM PMSF, pH 7.5). Protein concentrations were determined by absorbance at 280 nm and then TEV cleaved with 1:50 molar ration of TEV:Protein for 1 hour at room temperature. TEV cleaved mixture was added back to 5 mL bed volume of Talon resin to remove TEV and the HIS-MBP tag. Unbound fraction from the reverse IMAC was subjected to a Bradford assay to calculate protein concentration and flash frozen in LqN_2_ before -80 °C storage.

### Protein prep for Coq5p with and without the octapeptide

Protocol was essentially the same as above, except constructs containing truncated forms of Coq5p with and without the octapeptide were cloned and transformed. Also, all buffers contained 400 mM NaCl instead of 200 and pH was set to 7.8 instead of 8.0.

### Oct1p enzyme activity assay

Peptides were obtained from BIOsynthesis (see Key Resources table). Peptides arrived dry and were resuspended in 1x assay buffer (25 mM HEPES 100 mM NaCl pH=8.0) to a stock concentration of 60 μM. 600 pmol of peptide along with 40 pmol of Oct1p or Oct1p H558R was diluted to 100 μL of 1x assay buffer (25 mM HEPES 100 mM NaCl pH=8.0) in black-walled and round-bottomed plates. Florescence at an excitation wave length of 320 nm and emission wave length of 420 nm was measured for eight hours every five min. Slope calculations were made for each five-minute interval, and V_max_ was calculated using the 3^rd^ maximum slope observed between time points.

A no-enzyme control was performed alongside each peptide to assess peptide auto hydrolysis. The rate of auto hydrolysis (calculated as described for the enzymatic reactions) was subtracted from the V_max_ to obtain background subtracted enzyme activity rate. Averages and standard deviations of eight technical replicates were used in the calculations and reported.

Raw florescence values were converted to molar values with a proteinase K control. The same amount of peptide (600 pmol) was mixed with 4 pmol of proteinase K. Within 15 min, the proteinase K completely processed the 600 pmol of peptide. This florescence was assumed to be 100% conversion and the minimum florescence observed in the no enzyme control was assumed to be 0% converted. This standard was used to calibrate the fluorescence to pmol of peptide converted by Oct1p.

### Fluorescence microscopy

Yeast (1×10^8^ cells)transformed with various FLAG-tagged constructs were removed from cultures by pipetting and immediately fixed with formaldehyde (4% final concentration, gentle agitation on a nutator, 1 h, ~23 °C). The fixed cells were harvested by centrifugation (1000 *g*, 2 min, ~23 °C), washed three times with 0.1 M potassium phosphate pH 6.5 and once with K-Sorb buffer (5 mL, 1.2 M sorbitol, 0.1 M KPi, pH 6.5), and re-suspended in K-Sorb buffer (1 mL). An aliquot of the cells (0.5 mL) was mixed with K-Sorb-BME (0.5 mL, K-Sorb with 140 mM BME) and incubated (~5 min, ~23 °C). Zymolase 100T was added to 1 mg/mL final concentration and incubated (20 min, 30 °C). The resultant spheroplasts were harvested by centrifugation (1000 *g*, 2 min, ~23 °C), washed once with K-Sorb buffer (1.4 mL), and re-suspended in K-Sorb buffer (0.5 mL). A portion of the cells (0.25 mL) was pipetted onto a poly-D-lysine coated microscope coverslip and allowed to settle onto the slides (20 min, ~23 °C). To permeabilize the cells, the supernatant was aspirated from the coverslips, and MeOH (2 mL, –20 °C) was added immediately and incubated (6 min, on ice). The MeOH was aspirated and immediately replaced with acetone (2 mL, –20 °C) and incubated (30 s, on ice). The acetone was aspirated, and the slides were allowed to air-dry (~2 min). The samples were blocked with BSA (“BSA-PBS” [1% BSA in PBS], 2 mL, 30 min incubation at ~23 °C), and incubated with primary antibodies (Sigma F1804, 1 mg/mL stock anti-FLAG primary Ab at a 1:2000 dilution in PBS-BSA; in-house anti-Cit1p antibody (Guo et al., 2017) at a 1:500 dilution in PBS-BSA; 1 mL, 12 h, 4 °C). The samples were washed 5 times with PBS-BSA (2 mL, ~23 °C) and incubated with secondary antibodies diluted in PBS-BSA [1 μg/mL working concentration for each: Goat anti-Mouse IgG (H+L) Secondary Antibody, Alexa Fluor 594 conjugate (Thermo A-11005) and Goat anti-Rabbit IgG (H+L) Secondary Antibody, Alexa Fluor 488 (Thermo Cat# A-11008)] (1 mL, 2 h, ~23 °C, in the dark). The samples were washed 5 times with PBS-BSA (2 mL, ~23 °C) and twice with PBS (2 mL, ~23 °C). The last wash was aspirated and the slides were allowed to air dry briefly in the dark. The coverslips were mounted onto slides with 50% glycerol in PBS (8 μL) and imaged by fluorescence microscopy.

### Quantitative western blot for Coq5-FLAG constructs

WT yeast or Δ*oct1* yeast were transformed with Coq5p-FLAG or Coq5p F23A-FLAG. 3 individual colonies were selected from the transformation plate and grown overnight in –URA dropout media supplemented with 2% glucose. The following day, 2.5 × 10^6^ yeast were inoculated into 100 mL of –URA drop out media supplemented with 3% glycerol and 0.1% glucose. These were left to grow for a second night before 2.0×10^8^ cells were collected and processed for western blot. Yeast were lysed in 150 μL of lysis buffer (2 M NaOH, 1 M BME) for 10 min with periodic vortexing. Protein was TCA precipitated with 150 μL of 50% TCA and washed with 1 mL of Acetone. Protein pellet was resuspended in 120 μL of 0.1 M NaOH and 50 μL of 6x LDS sample buffer. 10 μL of this protein extract was run on a gel and subjected to Western blot analysis. Western data were quantified using Licor image studio. Ratios between Coq5p and actin were calculated and then averaged across the 3 biological replicates for each time point. These averages and standard deviations are reported.

### Growth curves for yeast

Single yeast were grown overnight in media with the proper selection +2% glucose. 5×10^6^ cells were collected from the overnight growth and spun down and resuspended in 1 mL of the growth curve media. 100 μL of this mixture was transferred to a clear round bottom sterile 96 well (eight individual replicates). The plate was covered with a breathable optical film and put inside the plate reader. The plate reader maintained temperature at 30°C during the duration of the run. The plate was constantly mixed and OD_600_ was measured every 5 min. Growth curve was constructed by taking the average and standard deviation of the eight replicates at a given time point.

### Targeted Q lipidomics

Single colonies of *S. cerevisiae* were inoculated into 5 mL URA drop out 2% glucose starter cultures and incubated for ~18 hr, 30 °C, 230 rpm. 100 mL URA drop out media(0.1% glucose, 3% glycerol) cultures were seeded with 2.5 x 10^6^ cells/mL and incubated at 30 °C, 230 rpm for 25 hr (OD between 1-2).Each treatment was done in biological triplicate. 5 x 10^8^ cells yeast were centrifuged (4 °C, 10 min, 3220 *g*) in 50 ml falcon tubes and cell pellets were flash frozen in N2(l). Yeast samples were thawed on ice and resuspended in 100 μL cold water. Samples were transferred to 2 mL tubes. To this, 100 μL glass beads (0.5 mm dia) were added and bead beat for 30 s (cold room). CoQ_10_ internal standard (10 μL, 10 μM) was added to each sample, followed by bead beating for 30 s. To this, 900 μL organic (1:1 CHCl_3_/MeOH, 4 °C) was added and vortexed (2 x 30 s). 33 μL 6 M HCl (4 °C) was added, followed by vortexing (2 x 30 s). Samples were spun (5,000 *g*, 2 min, 4 °C)to separate phases. After removing the aqueous layer, ~400 μL of organic was collected in a clean tube and dried under Ar_(g)_. Dried lipids were reconstituted in ACN/IPA/H_2_O (65:30:5, v/v/v, 100 μL) by vortexing (2 x 30 s, RT). Samples were transferred to a brown autosampler vial and stored under Ar before placing at -80°C

Ten microliters of lipid extract were injected by a Vanquish autosampler (Thermo Scientific) onto an Acquity CSH C18 column held at 50 °C (2.1 x 100 mm x 1.7 μm particle size; Waters). Mobile phase A consisted of 10 mM ammonium acetate in ACN/H2O (70:30, vol/vol) containing 250 μL/L acetic acid. Mobile phase B consisted of 10 mM ammonium acetate in IPA/ACN (90:10, vol/vol) with the same additives. Using a Vanquish Binary Pump (400 μL/min; Thermo Scientific) mobile phase B was held at 2% for 2 min and then increased to 30% over 3 min. Mobile phase B was then further increased to 85% over 14 min and then raised to 99% over 1 min and held for 7 min. The column was then reequilibrated for 5 min before the next injection. The LC system was coupled to a Q Exactive Focus mass spectrometer by a HESI II heated ESI source (Thermo Scientific). The MS was operated in positive and negative parallel reaction monitoring (PRM) mode acquiring scheduled, targeted PRM scans to quantify key CoQ intermediates. MS acquisition parameters were 17,500 resolving power, 1 × 106 automatic gain control (AGC) target for MS1 and 1 × 105 AGC target for MS2 scans, 25 units of sheath gas and 10 units of auxiliary gas, 300 °C HESI II and inlet capillary temperature.

#### Data Analysis

Peaks were automatically integrated using TraceFinder software (Thermo) and all integrations were checked manually.

### Half-life measurements for Coq5p

Four separate yeast colonies for both genotypes were grown overnight in 5 mL of synthetic complete media containing 2% glucose at 30 °C. The next day, 1.25×10^8^ cells were transferred into 500 mL of synthetic complete media containing 2% glucose and left to grow for a second night. The next morning, yeast was back diluted to an OD of 0.6 in 1 L of media and allowed to grow for 2 hours. At the 2 hour mark, 1 mL of 50 mg/ml cycloheximide stock in dimethyl sulfoxide was added the 1 L culture (50 μg/ml final). 2.0×10^8^ were collected at the time points indicated and added to 2x stop solution (20 mM NaN_3_ 0.5 mg/mL BSA) before being spun down and stored at -80°C

Yeast were lysed in 150 μL of lysis buffer (2 M NaOH, 1 M BME) for 10 min with periodic vortexing. Protein was TCA precipitated with 150 μL of 50% TCA and washed with 1 mL of Acetone. Protein pellet was resuspended in 120 μL of 0.1 M NaOH and 50 μL of 6x LDS sample buffer. 15 μL of protein was run on a gel and subjected to Western blot analysis.

Western data were quantified using Licor image studio. Ratios between Coq5p and VDAC were calculated and then normalized to the amount of protein at time 0. These normalized values were then averaged across the 4 biological replicates for each time point. These averages and standard deviations are reported. Half-life was calculated for each biological replicate, and then the average and standard deviation of the individually calculated half-lives were reported.

### Differential scanning fluorimetry

DSF was performed similarly to how it was previously described (Niesen et al., 2007). Coq5p protein prepared as described above. 300 pmol of Coq5p with or without the octapeptide on the end were diluted in to 20 μL final volume in DSF buffer (400 mM NaCl, 50 mM HEPES, 10% glycerol pH=7.8, final in reaction). A final concentration of 4x sypro orange (thermo S-6650) in the final reaction. Plate was loaded into a QuantStudio 6 quantitative thermo cycler. A temperature gradient from 4°C to 95°C over 1 hour. Sypro orange florescence was measured over the temperature gradient and data were analyzed in the Protein thermal shift software to assess melting temperature.

### QUANTIFICATION AND STATISTICAL ANALYSIS

See each individual method for the associated statistical analysis.

